# Regime shifts and hysteresis in the pitcher-plant microecosystem

**DOI:** 10.1101/087296

**Authors:** Matthew K. Lau, Benjamin Baiser, Amanda Northrop, Nicholas J. Gotelli, Aaron M. Ellison

## Abstract

Changes in environmental conditions can lead to rapid shifts in the state of an ecosystem (“regime shifts”), which, even after the environment has returned to previous conditions, subsequently recovers slowly to the previous state (“hysteresis”). Large spatial and temporal scales of dynamics, and the lack of frameworks linking observations to models, are challenges to understanding and predicting ecosystem responses to perturbations. The naturally-occurring microecosystem inside leaves of the northern pitcher plant (*Sarracenia purpurea*) exhibits oligotrophic and eutrophic states that can be induced by adding insect prey. Here, we further develop a model for simulating these dynamics, parameterize it using data from a prey addition experiment and conduct a sensitivity analysis to identify critical zones within the parameter space. Simulations illustrate that the microecosystem model displays regime shifts and hysteresis. Parallel results were observed in the plant itself after experimental enrichment with prey. Decomposition rate of prey was the main driver of system dynamics, including the time the system remains in an anoxic state and the rate of return to an oxygenated state. Biological oxygen demand influenced the shape of the system’s return trajectory. The combination of simulated results, sensitivity analysis and use of empirical results to parameterize the model more precisely demonstrates that the *Sarracenia* microecosystem model displays behaviors qualitatively similar to models of larger ecological systems.

## 1. Introduction

Regime shifts in ecological systems are defined as rapid changes in the spatial or temporal dynamics of an otherwise resilient system. Ecological regime shifts are caused by slow, directional changes in one or more underlying state variables, such as species abundance, dissolved oxygen content, or nutrients [1, 2]. Regime shifts are of particular concern when the return rate to a previous (and perhaps more desirable) state is slow or requires a larger input of energy or resources relative to what initiated the state change (i.e., hysteresis). In the last several years, many researchers have suggested that a wide range of ecological systems are poised to “tip” into new regimes [2, 3], or even that we are approaching a planetary tipping point [4]; but see [5]. Because identifying changes in the underlying state variables of most ecosystems require high frequency, long-term measurements [6], our understanding of the causes and consequences of ecological regime shifts has progressed relatively slowly. More rapid progress could be achieved by working with well-understood systems that can be described mathematically and manipulated experimentally over shorter time scales.

It is rare to find an ecological system in which the occurrence of a regime shift, and its cause-and-effect relationship with one or more underlying environmental drivers, is unambiguous [7]. This is primarily because long time series of observations collected at meaningfully large spatial scales are required to identify the environmental driver(s), its relationship to the response variable of interest, the stability of each state, the breakpoint between them, and hysteresis of the return time to the original state [3, 7]. Detailed modeling, and decades of observations, and experiments have led to a thorough understanding of one canonical example of an ecological regime shift: the rapid shift from oligotrophic (low nutrient) to eutrophic (high nutrient) states in lakes (e.g.,[8, 9]). The primary difficulties with using lakes as models for studying alternative states and ecological regime shifts are their large size (which precludes extensive replication:[10]) and the long time scales (decades) required to observe a regime shift, subsequent ecosystem hysteresis, and eventual recovery [11, 12]. Models of lake ecosystems and their food webs, and associated empirical data have revealed that recovery of lakes from a eutrophic to an oligotrophic state can be very slow—on the order of decades to centuries [12]—and depends not only on slowing or reversing directional changes in underlying state variables but also on the internal feedback dynamics of the system. Other aquatic systems, including fisheries [13], rocky intertidal communities, and coral reefs [3] have provided additional empirical support for these model results in terms of both dynamics and duration [14].

In a previous study, we experimentally demonstrated that organic-matter loading (i.e., the addition of excess insect prey to pitchers) can cause a shift from oligotrophic to eutrophic conditions in a naturally-occurring microecosystem: the water-filled leaves of the northern (or purple) pitcher plant, *Sarracenia purpurea* L. [15]. We use the term “microecosystem” here because the pitcher plant and its inquiline food web is a naturally occurring, co-evolved community of organisms, which is not necessarily the case for microcosms [16]. In the typically five-trophic level *Sarracenia* microecosystem, bacteria reproduce rapidly and drive the nutrient-cycling dynamics [17]. Prey additions cause shifts from oligotrophic to eutrophic states in hours or days rather than years or decades. Further, the comparatively small volume of individual pitchers, the ease of growing them in greenhouses and the occurrence of large, experimentally manipulable populations in the field [18] have allowed for replicated studies of trophic dynamics and regime shifts in a whole ecosystem.

Here, we build on the original mathematical model of the *Sarracenia* microecosystem [15], estimating parameter values using new empirical data and introducing more realism into the underlying environmental drivers of the model. We then use sensitivity analysis to identify the model parameters that most strongly control the dynamics of the system. We illustrate that once organic-matter input is stopped, the *Sarracenia* microecosystem—like large lakes—can eventually overcome the hysteresis in the system and return to an oligotrophic state. We conclude that the mathematical model illustrates dynamic behaviors that are qualitatively similar to models of regime shifts in lakes and other ecosystems, and we suggest that the *Sarracenia* microecosystem is useful model for studying ecological regime shifts in real time.

## 2. Methods

### 2.1. The pitcher-plant microecosystem

The eastern North American pitcher plants (*Sarracenia* spp.) are perennial carnivorous plants that grow in bogs, low nutrient (“poor”) fens, seepage swamps, and sandy out-wash plains [19]. Their leaves are modified into “pitchers” [20], tubular structures that attract and capture arthropods, and occasionally small vertebrate prey (e.g.,[21, 22]). In the pitchers, prey are shredded by obligate pitcher-inhabiting arthropods, including histiostomatid *Sarraceniopus* mites, and larvae of sarcophagid (*Fletcherimyia fletcheri*) and chironomid flies (*Metrocnemius knabi*) [23, 24, 25]. The shredded organic matter is further decomposed and mineralized by a diverse assemblage of microbes, including protozoa [26], yeasts [27], and bacteria [28].

Unlike other species of *Sarracenia* that also secrete and use digestive enzymes to extract nutrients from their captured prey, *S. purpurea* pitchers secrete digestive enzymes for only a fraction of their lifespan [29]. Instead, *S. purpurea* relies on its aquatic food web to decompose the prey and mineralize their nutrients [30]. As a result, the rainwater-filled pitchers of *S. purpurea* are best considered a detrital-based, “brown” ecosystems in which bacterially-mediated nutrient cycling determines whether it is in an oligotrophic or eutrophic state [15, 17, 31].

### 2.2. Oxygen dynamics in lakes and pitchers

Oxygen dynamics, in both lakes and *Sarracenia* pitchers, can be described using a simple model that yields alternative oligotrophic and eutrophic states and hysteresis in the shift between them [1]:

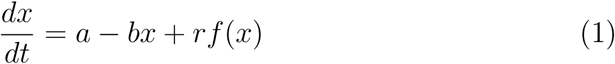

In this model, the observed variable *x* (e.g., oxygen concentration) is positively correlated with state variable *a* (e.g., rate of nutrient input or photosynthesis), and negatively correlated with state variable *b* (e.g., rate of nutrient removal or respiration). The function *r f* (*x*) defines a positive feedback that increases *x* (e.g., the rate of nutrient recycling between the sediment in lakes or mineralization-immobilization by bacteria of shredded prey in a water-filled *Sarracenia* pitcher). If *r* > 0 and the maximum of {*r f* (*x*)} > *b*, there will be more than one equilibrium point (i.e., stable state) [1]; the function *f* (*x*) determines the shape of the switch between the states and the degree of hysteresis.

Following [1], we used a Hill function for *f* (*x*):

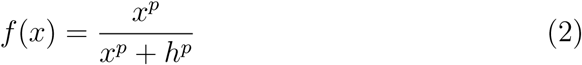

The Hill function provides a simple model that can produce threshold behaviors. The dynamics of the state variable *x* is determined by parameters *p* and *h*, which determine the rate of change and the in flection point of the curve, respectively (Fig. 1A). If *p* is set such that more than one possible state exists for the system, *h* determines the threshold for the transition between these states. When viewed in phase-space (Fig. 1B), the transition between states can be seen as a path traversed by the system between distinct regions (i.e., phases). In part because of this threshold property, the Hill function has been applied to systems ranging from biochemistry and microbiology to ecology, whose dynamics depend on a limiting resource (e.g.,[32].

**Figure 1:**
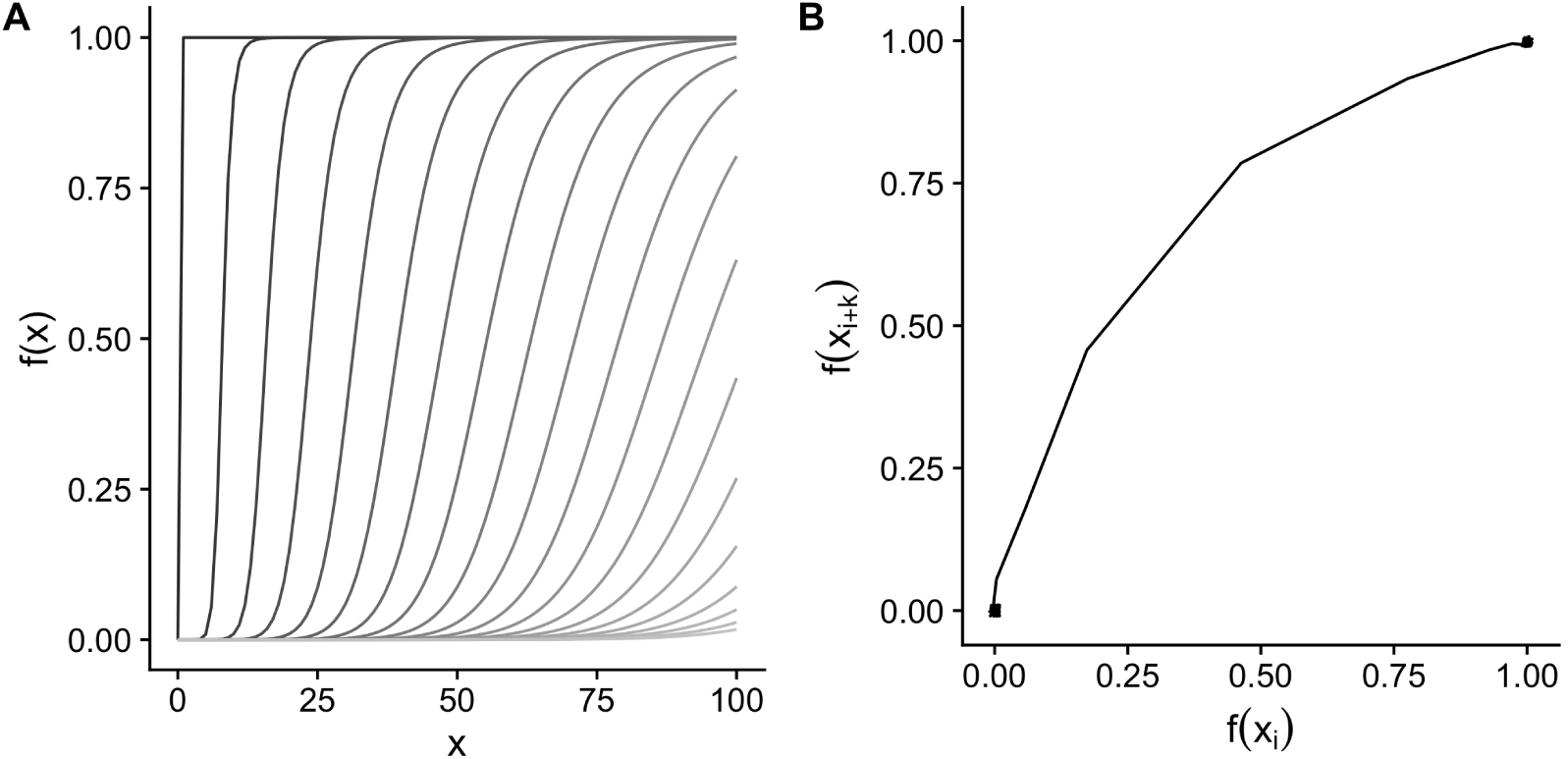
The threshold dynamics of the Hill function are determined in part by the in flection parameter *h*. A) Plotted output of the Hill function for different values of *h* (different lines shaded darker for lower values), ranging from 0.1 to 150 with *p* = 10. B) Lagged (*k* = 1 lag term) phase plot of the Hill function with *h* = 71.11, showing the state transition (lower-left to upper-right). A small amount of random variation was introduced to the series to reveal overlapping points within the two states.

We modeled the dynamics of the trophic state of the *Sarracenia* microecosystem using an equation of the same underlying form as Eq. 1:

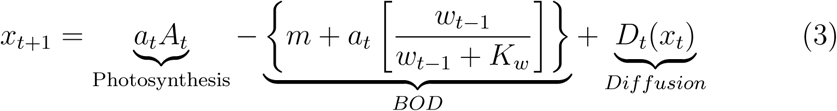

Each model term is described below and summarized in Table 1.

**Table 1:** Terms, units, and interpretation for the model of oxygen dynamics in pitcher plant fluid.

| Term | Units | Interpretation |
| --- | --- | --- |
| $t$ | minutes | Time (model iteration) |
| $f$ | $1/t$ | Constant adjusting sine wave of diurnal PAR for frequency of measurements |
| $x_t$ | $\text{mg L}^{-1}$ | [O<sub>2</sub>]: concentration of oxygen in the pitcher fluid |
| **Photosynthesis** |  |  |
| $A_t$ | $\text{mg L}^{-1}$ | Production of oxygen by photosynthesis, and infused from the plant into the fluid during the day |
| $a_t$ | $\text{mg L}^{-1}$ | Photosynthetic rate augmentation by microbial nutrient mineralization |
| $a_{min}, a_{max}$ | $\text{mg L}^{-1}$ | Minimum or maximum possible photosynthetic augmentation |
| PAR | $\mu\text{mol} \cdot \text{m}^{-2} \cdot \text{s}^{-1}$ | Photosynthetically active radiation |
| **Respiration** |  |  |
| $w_t$ | mg | Mass of prey remaining at time $t$ |
| $W$ | mg | Maximum mass of decomposable prey (set at $75\mu\text{g}$) |
| $K_w$ | $\text{mg min}^{-1}$ | Half-saturation constant for bacterial carrying capacity |
| $m$ | $\text{mg L}^{-1}$ | Basal metabolic oxygen used by bacteria (respiration) |
| **Nutrients** |  |  |
| $n_t$ | $\text{mg L}^{-1}$ | Quantity of nutrients mineralized by decomposition; a function of $w_t$, and $x_t$ |
| $s$ | dimensionless | Sigmoidal curve steepness relating nutrient mineralization to augmentation |
| $d$ | mg | Inflection point of sigmoidal curve relating mineralization to augmentation |
| $\beta$ | dimensionless | The rate of prey decomposition |

The model (Eq. 3) of the *Sarracenia* microecosystem (Fig. 2A) is made up of the two main terms: production of oxygen by photosynthesis, and use of oxygen (respiration) during decomposition (BOD: biological oxygen demand). The pitcher fluid is oxygenated (*x*) at each discrete time step (*t*, in minutes) as the plant photosynthesizes (*A_t_*). The value of *A_t_* is determined by sunlight, which we modeled as a truncated sine function producing photosynthetically active radiation (PAR) (Fig. 2B), and by the maximum photosynthetic rate (*A_max_*) (Fig. 2C), which leads to the production of dissolved oxygen in the pitcher fluid (Fig. 2D). Photoautotrophic bacteria, but not algae, have been detected in a recent molecular study of the proteome of *S. purpurea* inquiline communities; however, the total number of peptides mapping to these bacterial species was low relative to other species in the community [33]. Thus, although our model does not explicitly account for other photosynthetic organisms, these previous findings suggest that their contributions to the oxygen dynamics of the pitcher-plant fluid are likely to be small relative to that of the pitcher plant itself.

**Figure 2:**
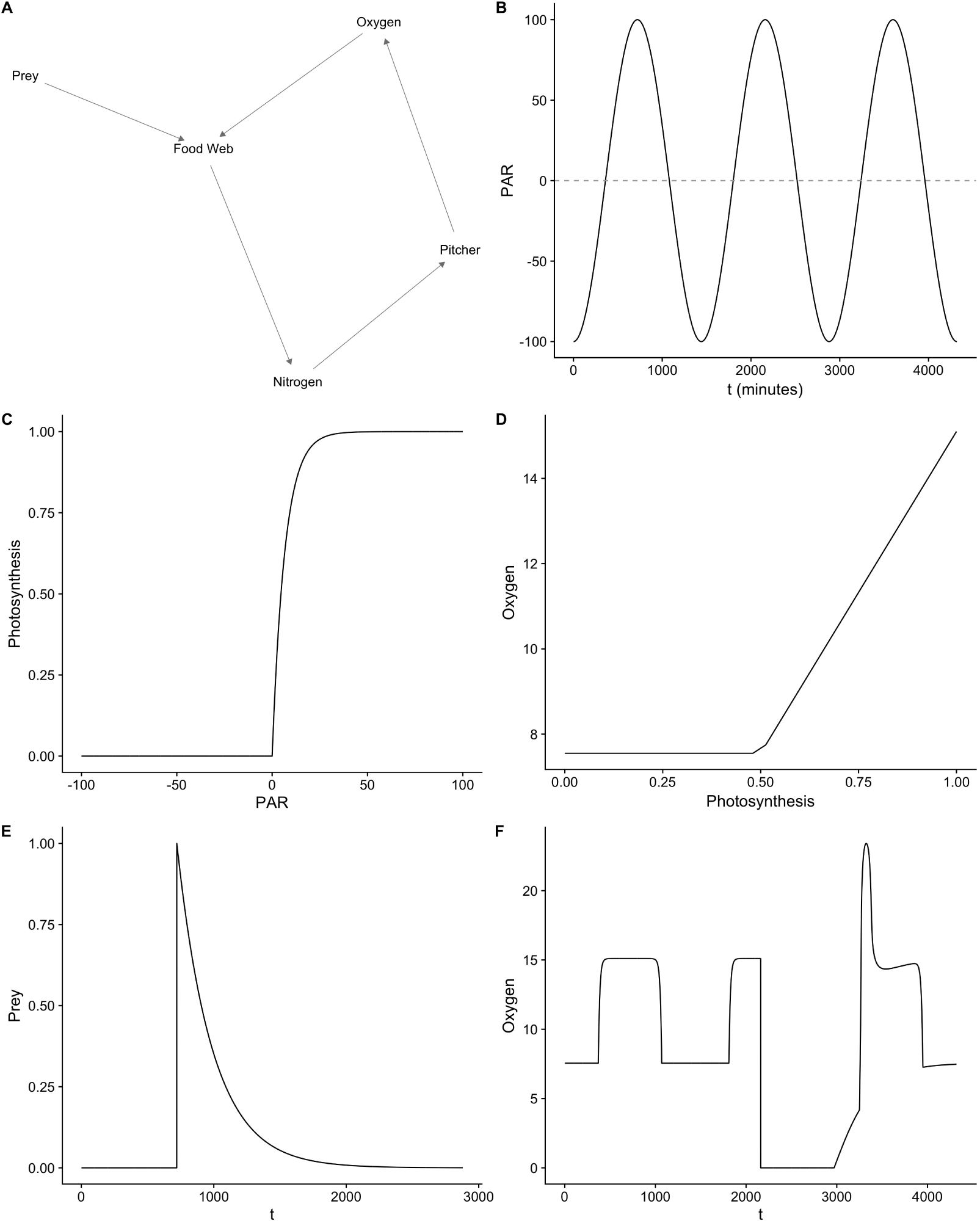
A) The pitcher plant model shown as a network diagram. The nodes in the graph and their corresponding variables in the model are Prey (*w*), the microbially-dominated food web (controlled by *K_w_*), Nitrogen (*n*), Oxygen (*x*) and the pitcher plant itself, which is included to show the fluxes of nitrogen and oxygen as they relate to the plant. B) Photosynthetically active radiation (PAR) from the sun or other light source modeled as a sine wave. Although negative values of PAR are set equal to zero in our model’s photosynthesis function, we show here the full sine-wave to illustrate the mathematical function from which PAR is derived. C) The relationship between PAR and photosynthesis in the pitcher plant. D) The output of dissolved oxygen in the pitcher fluid as a function of pitcher-plant photosynthesis. E) The decomposition of prey over time. F) The impact of prey addition (*t* = 2160 min) on dissolved oxygen in the pitcher-plant fluid.

Decomposition of shredded prey by bacteria requires oxygen. The oxygen demand from respiration is modeled by the BOD term in Eq. 3. The parameter *m* is the basal metabolic respiration of the food web with no prey to decompose in the system. Adding prey (*w*) induces decomposition, which we model as a negative exponential function with rate parameter *β*, and a constant *W* (maximum prey mass decomposed over 48 hours) using Eq. 4, and illustrated in Figure 2E.

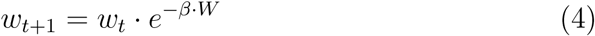

Bacterial populations increase at a rate determined by a half-saturation function with parameter *K_w_* (Eq. 3), which increases BOD, and the depletion of oxygen from the pitcher fluid (Fig. 2F). Demand of the food web for oxygen (i.e., BOD) depends on the decomposition rate (*β*) and the shape parameter (*K_w_*), but only when prey is present in the system (*w*_*t*−1_ > 0 in Eq. 3). When prey is absent (i.e., *w*_*t*−1_ = 0), BOD terms simplify by multiplication to the basal metabolic rate (*m*).

Photosynthesis may be limited by available nutrients, primarily nitrogen and phosphorus [34, 35], that are mineralized by bacteria from the prey [17]. Photosynthesis is augmented (*a_t_*) by nutrient mineralization rate (*s*). We model *a_t_* as a saturating function with bounds determined by the range terms (*a_min_*, and *a_max_*), *s*, and the point of saturation (*d*):

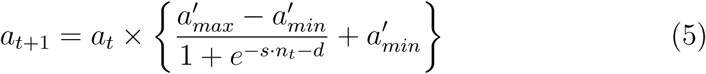

The other impact of prey addition, and subsequent decomposition by the food web, is the release of nutrients into the pitcher fluid. The mineralization variable *n_t_* (Eq. 5), which is modeled as proportional to the product of the amount of oxygen and prey in the system (i.e., *n_t_*_+1_ = *c* · (*w_t_* · *x_t_*) where *c* is a constant of proportionality), creates a feedback from decomposition to oxygen production by the plant (i.e., the path in Fig. 2A from the food web to nutrients to pitcher to oxygen and back to the food web).

Thus, the mineralization term couples respiration (oxygen depletion) to photosynthesis (oxygen production) when prey is introduced to the system, and the food web begins to decompose the prey and release nutrients into the pitcher fluid. Finally, a small amount of oxygen diffuses into the pitcher fluid directly from the atmosphere (*D*(*x*)); however, the diffusion rate is generally so low that it is negligible relative to changes resulting from photosynthesis and BOD.

### 2.3. Estimating decomposition rate

We used data from a greenhouse pitcher-feeding experiment to estimate the decomposition parameter. The experiment was conducted over 35 days, starting on July 6, 2015, in a temperature-controlled greenhouse at the University of Vermont’s Biological Research Complex (Burlington, Vermont, USA). Eighteen newly-formed pitchers > 8 ml in volume were rinsed with deionized water, and randomly assigned to one of three organic matter addition treatments: 0.0, 0.5, or 5.0 mg mL^−1^ of pitcher fluid. Pitcher fluid was collected from randomly-selected pitchers at Molly Bog (Morristown, Vermont, USA: 44.500 N,-72.642 W) on the morning of 6 July 2015. The fluid was transported to the greenhouse, filtered through the 30-micron frit bed of a chromatography column (BioRad, Hercules, California, USA) to remove larger organisms, homogenized and added to experimental pitchers in the greenhouse.

Pitchers were loaded with a single pulse of organic matter every morning for the first four days. In this experiment (and following [36]), the organic matter was bovine serum albumin (BSA), DNA from salmon testes at a concentration of 1.5 *µ*g L^−1^ of BSA, and trace elements, potassium, calcium, sodium, magnesium and manganese in a ratio of 1:0.115:0.044:0.026:0.0001. In a pilot study, BSA yielded similar changes in dissolved oxygen as we had obtained previously using ground wasps as organic matter [15], and its use enabled us to measure changes in organic matter content via a simple Bradford assay, and in a pilot study, yielded similar changes in dissolved oxygen as we had obtained previously using ground wasps as organic matter (Sirota et al. 2013).

Three 100 *µ*L aliquots of pitcher fluid were sampled from single pitchers of each of 14 individual plants. Sampling was conducted twice a day from day 0 to day 20 at 8:30am and 5:00pm (± 2 hrs), once per day from day 20 to day 28 at 8:30am (± 2 hrs), and once each on days 30, 31, 33, and 35 (8:30am ± 2 hrs). Prior to sampling, the pitcher fluid was stirred with the pipette submerged to get fluid representative of an average of the entire pitcher. During sampling care was taken to minimize the introduction of additional oxygen from the sampling process itself. Pitcher fluid was topped off with deionized water after sampling to keep initial pitcher volumes consistent over the course of the 35 days. The initial volumes varied from 8 mL to 28.6 mL with a mean of 16.7 mL and standard deviation of 4.82 mL. Although we did control for the water taken with each sample, we did not adjust for the removal of nutrients and microscopic organisms that occurred at each sampling.

Samples were centrifuged at 13000*g*, after which the supernatant containing soluble BSA was removed, placed in a sterile tube and frozen at −80 °C until analyzed. A simple Bradford assay (Bradford 1976) was used to determine the concentration of BSA in each of the pitcher fluid samples. The assay was done using Bradford reagent (VWR), and the absorbance of each sample was measured on a Biophotometer Plus spectrophotometer (Eppendorf) at an optical density of 600 nm. Samples were read randomly on the spectrophotometer to avoid reaction time as a confounding variable. Sample concentrations were determined by comparison to standard curves generated using R [37].

We used an empirical least-squares estimator (LSE) approach to generate the best-fitting values for the decomposition parameter (*β*) in Eq. 4, given the quantity of prey added in the experiment and the duration of the prey addition. As *K_w_* is not a part of the decomposition function, we did not vary it during parameter estimation and model fitting. We then ran a series of 35-day simulations (equivalent to the run-time of the prey-addition experiment) in which *β* was sampled from a grid of values ranging from 1E-8 to 0.0007, and the amount of prey in the simulation was recorded at each simulated minute. For each run, the sum of squared errors (SSE) was recorded as Σ(*sim* − *obs*)^2^. The *β* that minimized the SSE in each simulation was considered to be the best-fit value for each replicate pitcher (*n* = 12). All model fitting was also done using R [37].

### 2.4. Sensitivity Analysis

We used a sensitivity analysis in which we varied the prey addition rate (*w*), decomposition rate (*β*), and the half-saturation constant *K_w_* to explore the behavior of the microecosystem model across a wide range of parameter space. Rather than set combinations of fixed values for the three parameters of interest, we sampled the model parameter space by drawing values independently from uniform distributions: *K_w_* ~ U(0.0001, 3), *β* ~ U(1.0E-6, 2.0E-3) and *w* ~ U(0, 100). To characterize baseline (oligotrophic) oxygen concentrations, for each combination of *β* and *K_w_* we ran one simulation in which no prey was added to the system (*w* = 0). In all simulations (*n* = 15,000) variables were initialized to 0 with the exception of oxygen (*x*), which was initialized using the photosynthesis term (*x*_0_ = 7.55). Simulated prey additions occurred at mid-day on days 4–6 (i.e., *t* = 6480 to *t* = 9360), each simulation ran for 30 simulated days (43,200 minutes), and output was saved for each simulated minute. The simulations were initialized using a random sample of parameter values, and run in parallel. Because the model is completely deterministic, the resulting runs can be reproduced by starting the simulations with the exact values used to initialize and parameterize the models.

To aid in the detection of the impact of the most important parameters and variables, in all simulations we set some parameter values to zero, which altered the model in the following two ways. First, we ignored the *D*(*x_t_*) term because we assumed that the amount of oxygen diffusing directly into the pitcher fluid from the atmosphere would be orders of magnitude lower than oxygen produced by pitcher photosynthesis [38]. Second, we noted that since the basal metabolic respiration of the food web parameter (*m*) is an additive constant, any change in the value of the constant *m*, (basal respiration of the microbial community) would result only in a proportional change in value of *x*, not in the shape (i.e., *dx/dt*), of the oxygen production over time. Thus, we could set *m* = 0 without loss of generality.

By setting *m* = 0, we also observed that the photosynthetic augmentation term (*a_t_*) in fluenced photosynthesis (*A_t_*), and BOD 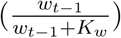 identically. Therefore, the parameters *s* and *d* in Eq. 5 could be set as constants in the sensitivity analysis. By ignoring diffusion, setting *m* = 0, and fixing *s* and *d*, we reduced the dimensionality of the sensitivity analysis to three (*w, β*, and *K_w_*), which eased the interpretation of the results.

We calculated two measures of the state of the system from the time series of oxygen concentration (*x_t_*): hypoxia, and return rate. We defined hypoxia in the model to be an oxygen concentration of ≤ 1.6 mg L^−1^, which is the median lethal O_2_ concentration ([O_2_]) for aquatic animals [39]. We measured the return rate of the system as the linear trend in [O_2_] (i.e., after removing the daily cycle in oxygen resulting from photosynthesis) across the entire simulation using Pearson’s correlation coefficient. Although the return trajectory can be non-linear, the linear trend measures the gross trends of returning (positive), little impact of feeding (zero) or remaining at depressed oxygen levels.

### 2.5. Code Availability, and Execution

The model was coded in the **R** programming language [37]. The 15,000 model runs for the sensitivity analysis were run on the Odyssey Supercomputer Cluster at Harvard University (Research Computing Group, FAS Division of Science, Cambridge, Massachusetts). Data, code for the simulations, and output of analyses are available in the Harvard Forest Data Archive (harvardforest.fas.harvard.edu/harvard-forest-data-archive).

## 3. Results

The equation representing decomposition and BOD resembles the Hill function in a general model of state changes with hysteresis (Eqs. 1 & 2). In general, when a Hill function is used in a basic alternative states model (e.g., *r f* (*x*) > *b* in Eq. 1), the inflection point (e.g., half-saturation constant *K_w_*) determines the threshold (Fig. 1A). Thus, modeling decomposition and BOD using a Hill function provided us with sufficient flexibility to yield a variety of state changes.

The simulations with the model produced dynamics observed in the empirical pitcher plant microecosystem. Because photosynthesis is nutrient-limited in *Sarracenia* [35], addition of prey increased modeled photosynthesis (Fig. 3A) relative to oligotrophic, prey-free, pitchers. In the oligotrophic state, and when no prey was added, BOD remained low throughout the entire simulation (black line in Fig. 3B). After prey was added on, for example, days 4–6 (*t* = 6480 to *t* = 9360 minutes), the system jumped into its alternative state: BOD increased rapidly then declined slowly as prey was mineralized (grey line in Fig. 3B). The combination of the smooth, slow recovery response of photosynthesis to prey addition and the abrupt shift in BOD following prey addition (Fig. 3A & B) resulted in an abrupt shift in the system from an oxygenated state into an anoxic state and a very slow (hysteretic) recovery (Fig. 3C). The hysteresis of the system was apparent when oxygen concentration was plotted as a time-lagged phase plot (lag = 1440 minutes starting at *t*=720), which shows the change in oxygen following addition of prey at *t* = 6480 and the slow return due to high BOD (Fig. 3D). These results were corroborated by observations from field and greenhouse experiments in which oxygen was observed to decline with the capture or addition of insect prey to the pitcher [15], and demonstrate the presence of both state changes and hysteresis (i.e., Fig. 3D) for at least some parameterizations of the model.

**Figure 3:**
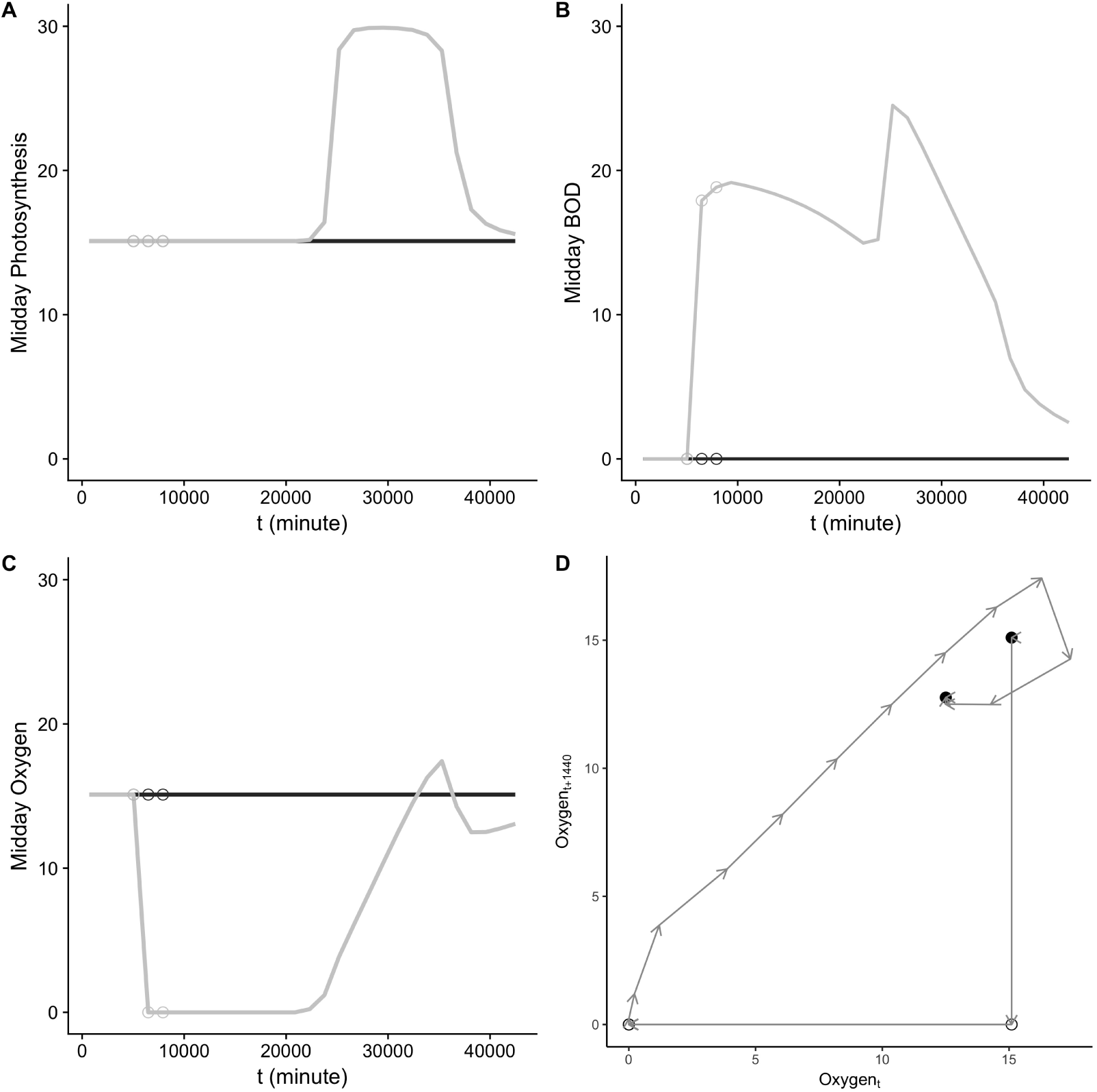
In model simulations, addition of prey impacts both photosynthetic oxygen production via augmentation from nutrients mineralized from prey and oxygen depletion through the biological oxygen demand (BOD) of microbial metabolism. (A) shows how photosynthesis increases when prey is added (grey) on days 4–6 (*t* = 5040 to *t* = 7920 minutes; indicated by open circles), relative to when no prey was added (black). (B) The quantity of oxygen used via the BOD of microbial decomposition. The net impact in this parameterization was a decrease in dissolved oxygen when prey was added to the system; (C) shows oxygen present in the pitcher at mid-day. (D) A time-lagged phase plot (*t*_0_ = 720, lag = 1440 min) showing the change in oxygen production during the prey addition simulation. Beginning and end points of the simulation are indicated by closed circles. When prey was added at *t* = 5040, *t* = 6480, and *t* = 7920 (open circles), it was decomposed rapidly by the microbially-dominated food web, resulting in oxygen depletion. The altered return trajectory (i.e., hysteresis) resulting from the biological oxygen demand in the system is shown by the arrows indicating the direction of the change in oxygen through time.

The parameter fitting and sensitivity analysis revealed several key effects of the parameters that we varied. First, the LSE model-fitting procedure resulted in an estimate for *β* of *x̅* = 0.00041 ± 0.0004 [se] (Fig. 4, vertical lines). Second, varying *β* had a large effect both on the percent time spent in an hypoxic state and on the return rate (steeper contours with increasing *β* in Fig. 4). Last, varying the amount of prey by two orders of magnitude produced a sharp threshold for the effect of varying *β* on hypoxia and return rate (Fig. 4).

**Figure 4:**
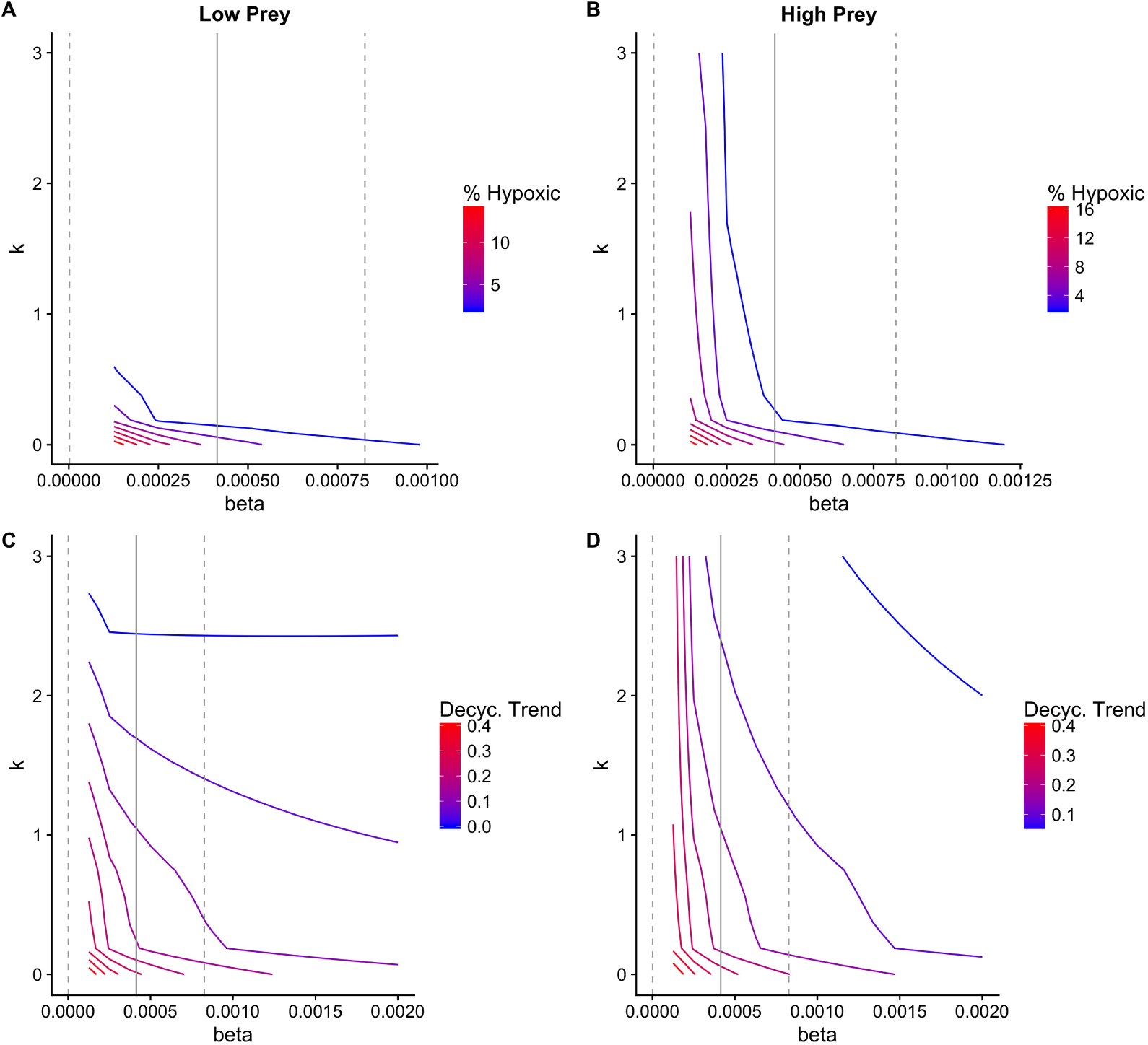
Sensitivity analysis of the pitcher-plant model revealed non-linear effects of varying the parameters *β* and *K_w_* (*n* = 15000 simulations). Contour plots show the percent time the system spent in an hypoxic state (top row), and the Pearson correlation coeffcient for the decycled trend (bottom row). The sensitivity simulations were repeated for additions of prey corresponding to 1 mg mL^−1^ (left column), and 100 mg mL^−1^ (right column) of prey added to the microecosystem. The LSE estimate for *β* (± 1 se) is plotted in each contour plot as vertical solid and dashed lines, respectively.

Although varying *β* has a potentially larger effect on the dynamics of the microecosystem than varying *K_w_*, the latter played an important role in determining the return trajectory of the oxygen. For simulations with lower values of *K_w_*, the oxygen concentration was still exponentially increasing when the simulation ended (Fig. 5A). Relative to simulations with higher *K_w_*, the return rate was faster when *β* was low enough, and there was prey (i.e., *w_t_*) remaining in the pitcher at the last observed time (Fig. 5B). Thus, in this part of the parameter space, if another round of feeding were to occur at a similar level of prey input, the system would never recover, and would remain in or near an hypoxic state.

**Figure 5:**
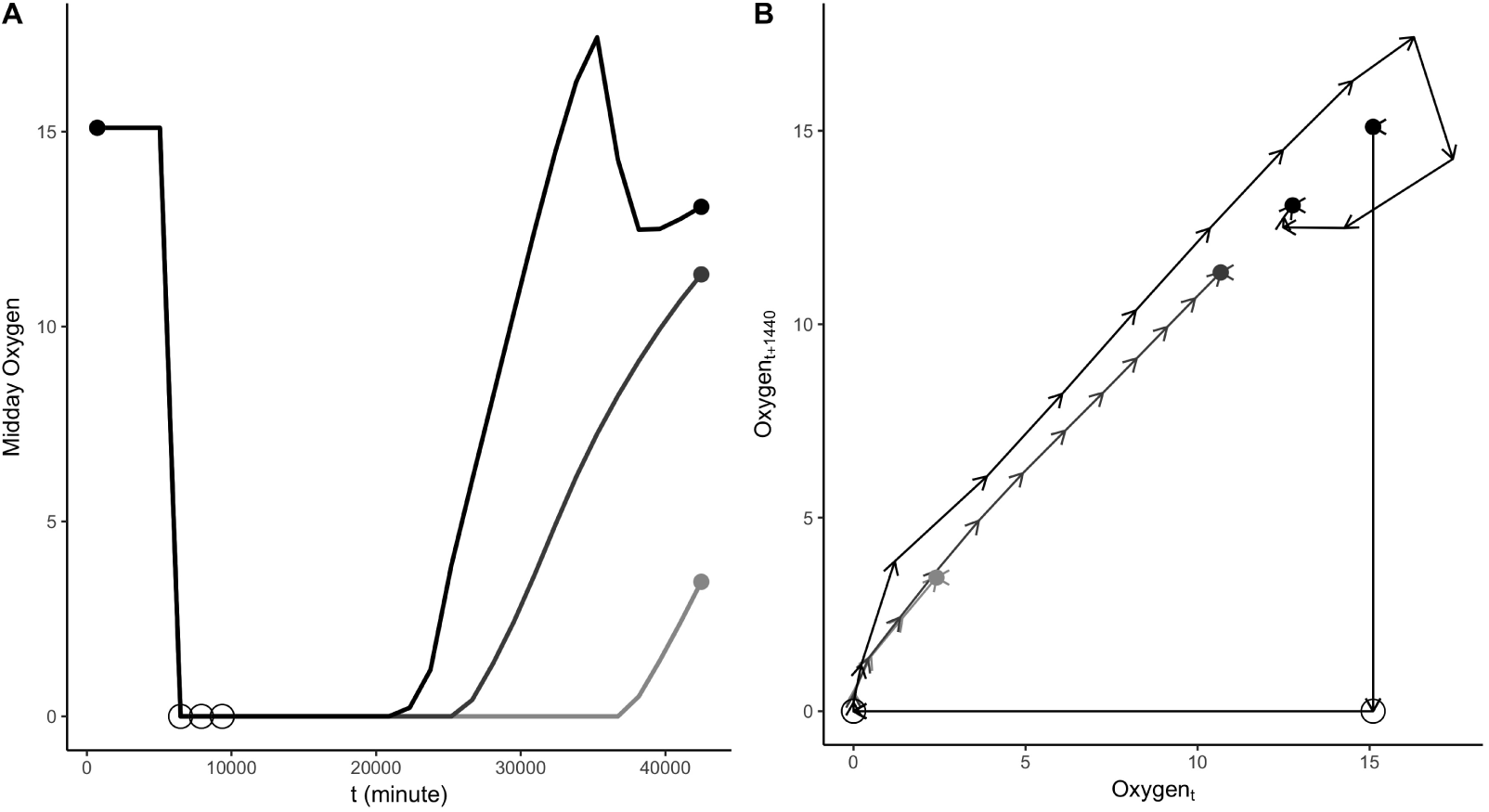
Oxygen dynamics in three simulations using different levels of bacterial carrying capacity (*K_w_*; light-grey = 0.1, dark-grey = 0.5, and black = 1) with the same rate of decomposition (*β* = 2.0E-6) illustrating hysteresis (i.e., altered return trajectory) of the system. (A) Lower levels of *K_w_* produce slower return rates over the course of the simulation. Prey addition (open circles) depressed mid-day oxygen curves at lower values of *K_w_*. Closed circles indicate the first and last mid-day prey addition points. (B) A time-lagged (*t* = 1440 minutes) phase plot for the same simulations showing that lower values of *K_w_* led to the oxygen being at lower levels for more time following prey addition (open circles), but followed a similar return trajectory as prey was decomposed by the food web (closed circles also indicate the beginning and end of each series). Although all three simulations ran for the same amount of time, the lengths of the trajectories are different in phase space because lower values of *K_w_* resulted in the system spending more time with the same amount of oxygen (i.e., *x_t_* = *x*_*t*+1440_).

## 4. Discussion

General theoretical work in complex systems has suggested that the definition of system boundaries is arbitrary and carries the potential for system dynamics to be mechanistically connected to, but unpredictable from, lower levels (or scales) of organization [40, 41]. However, others have argued that food-web dynamics of whole ecosystems can be inferred from the components (i.e., motifs and modules) of these ecosystems [42]. Overall, our model of the *Sarracenia* microecosystem supports the latter assertion: a focus on particular pathways (e.g., photosynthesis, decomposition, etc.) reproduced the non-linear behavior of its oxygen dynamics, including state changes and hysteresis. The results of the sensitivity analysis also revealed that the carrying capacity of the bacterial community (as simulated by the effect of *K_w_*) could contribute to observed non-linear state-changes of the *Sarracenia* microecosystem.

Predictions based on the model are highly sensitive to changes in the parameterization of decomposition (e.g. *β*). In the initial parameterization of this model, we started with an empirical estimate of decomposition rate in which > 99% of the average amount of prey captured could be decomposed in a single day [15, 43]. This is extremely rapid decomposition relative to a set of 58 previously published food webs [44], in which 1.27–66.2% of available detritus or organic matter is decomposed each day. When we set the decomposition parameter (*β*) equal to 2.57E-6, the overall decomposition rate approached the mean of the published food webs (24.22 ± 2.79% [SE]). This value for *β* is within the parameter space that we observed experimentally, and used in our sensitivity analysis, and suggests that insights gained from the *Sarracenia* microecosystem should be scalable to larger systems.

Although the dynamics of the *Sarracenia* microecosystem share similarities with lake, stream and other large-scale ecosystems, there are several differences that should be noted. First, oxygen levels in the pitcher plant are dynamically controlled by photosynthesis of the plant that serves as a strong driver of oxygen levels. In lakes, the primary oxygen production is carried out by phytoplankton, which are immersed in the aquatic system. Second, lake food webs are “green” (i.e., plant-based); whereas pitcher plant food webs are “brown” (detritus-based [17]). In lakes, the shift to a eutrophic state occurs through addition of limiting nutrients (usually N or P), accumulation of producer biomass that is uncontrolled by herbivores (see [45], and subsequent decomposition that increases biological oxygen demand [46, 47]. The *Sarracenia* microecosystem’s “brown” food web also experiences an increase in oxygen demand and microbial activity; however, this occurs during the breakdown of detritus that is characteristic of its shift from an oligotrophic to a eutrophic state [15]. Even though the source of the nutrients in the *Sarracenia* microecosystem is “brown”, the functional shape of the pathways involved in its nutrient cycling are similar to those in lakes with “green” food webs and are likely to lead to similar qualitative dynamics of both systems.

The results of our model and sensitivity analyses, combined with previously published empirical data [15], suggest that the *Sarracenia* microecosystem could be employed as a powerful system with which to develop new understanding of the dynamics of complex ecosystems. The food web of *S. purpurea* consists of species that share an evolutionary history, multiple trophic levels, and interactions that have been shaped by both environmental and co-evolutionary forces [21, 48]. Its abiotic environment and food web are comparable in complexity to large lakes [18, 49, 50]. It features similar critical transitions and non-linear dynamical behavior that are of broad interest for theoretical ecologists.

Mesocosm studies have been critiqued for lacking any or all of these characteristics [10], but a recent meta-analysis of the scaling relationships of the half-saturation constant (*K_w_*) provides evidence that uptake of nutrients such as nitrogen and phosphorus by food webs, and inter-trophic nutrient transfers, all are nearly invariant to spatial scale [32]. At the same time, the dynamics of the *Sarracenia* microecosystem play out over days, rather than years, decades, centuries, or even longer. Thus we conclude that, similar to previous work that has demonstrated the ability to scale up ecosystems processes (e.g., [51]), the pitcher-plant microecosystem provides an experimental system and computational model with which to study the linkages between “green”, and “brown” food webs [17, 52, 53] in the context of a food-web with an evolutionary history. Therefore, this system provides a powerful tool for identifying early warning signals of state changes in ecosystems that are of crucial importance for environmental management [54, 55].

## 5. Acknowledgments

We would like to thank Drs. Judith Bronstein and Greg Dwyer, and four anonymous reviewers for constructive feedback that greatly improved the manuscript. This work was made possible by grants from the National Science Foundation, including DEB 11-44056 and ACI 14-50277.

